# Rapid mini-chromosome divergence among fungal isolates causing wheat blast outbreaks in Bangladesh and Zambia

**DOI:** 10.1101/2022.06.18.496690

**Authors:** Sanzhen Liu, Guifang Lin, Sowmya R. Ramachandran, Giovana Cruppe, David Cook, Kerry F. Pedley, Barbara Valent

**Affiliations:** Department of Plant Pathology, Kansas State University, Manhattan, KS 66506-5502, USA; Foreign Disease-Weed Science Research Unit, United States Department of Agriculture–Agricultural Research Service (USDA-ARS), Ft. Detrick, MD 21702-5023, USA

**Keywords:** wheat blast, mini-chromosome, *Magnaporthe*, effector

## Abstract

Global wheat production is seriously threatened by the filamentous fungal pathogen, *Magnaporthe oryzae*, causing wheat blast disease. The pathogen was first identified in South America and recently spread across continents to Bangladesh (South Asia) and Zambia (South-central Africa). *M. oryzae* strains closely related with a South American field isolate B71 was found to have caused the wheat blast outbreaks in South Asia and Africa. Here, we studied the genetic relationship among isolates found on the three continents. Using an improved reference genome for B71 and whole genome sequences of isolates from Bangladesh, Zambia, and South America, we found strong evidence to support that the outbreaks in Bangladesh and Zambia were caused by the introductions of genetically separated isolates. Structural variation analysis using whole genome short-read sequencing data indicate all isolates closely related to B71 maintained at least one supernumerary mini-chromosome and, interestingly, some Zambian isolates contain more than one mini-chromosome. Long-read sequencing and *de novo* genome assemblies of two Zambian isolates show that both contain a mini-chromosome similar to the B71 mini-chromosome, although pervasive structural variation exists among them. Genome assemblies also provide evidence that one Zambian isolate carries an additional mini-chromosome that is highly divergent from the B71 mini-chromosome. Our findings show that while the core genomes of the multiple introductions are highly similar, the mini-chromosomes have undergone marked diversification. The maintenance of the mini-chromosome during the multiple introductions, and the rapid sequence and structural variation suggests the mini-chromosomes may serve important virulence or niche adaptation roles under diverse environmental conditions.

## INTRODUCTION

*Magnaporthe oryzae* (synonym of *Pyricularia oryzae*) is the ascomycetous fungus that causes blast diseases on a wide variety of grass species, including important grain crops (Gladieux *et al*. 2018; Valent *et al*. 2020; Ristaino *et al*. 2021). The *M. oryzae Oryza* pathotype (MoO), the lineage causing blast disease on rice (*Oryza sativa*), is reported to have arisen via host shift from a *Setaria* population around the time of rice domestication in China ∼7000 years ago (Couch *et al*. 2005). MoO subsequently dispersed world-wide through movement in infected seed, either through human movement or seed exchange (Saleh *et al*. 2014). In contrast, wheat blast disease caused by the *M. oryzae Triticum* pathotype (MoT) emerged in Brazil in 1985 and has spread within South America for less than four decades (Cruz and Valent 2017; Ceresini *et al*. 2018). Wheat blast jumped continents for the first time to Bangladesh in South Asia in 2016 (Malaker *et al*. 2016; Islam *et al*. 2016), presumably through importation of contaminated wheat seed/grain from South America (Ceresini *et al*. 2018). The disease is now established in Bangladesh, posing a threat to food security in Asia, including India and China, two major wheat producing and consuming countries. In 2017, wheat blast was identified in Zambia, Africa (Tembo *et al*. 2020), sounding the alarm of the world-wide spread of this recently-emerged disease.

Phylogenetic analyses of more than 2,500 genes and of whole-genome SNP data showed that MoT isolates from Bangladesh were genetically close to Bolivian MoT isolate B71 collected in 2012 (Gladieux *et al*. 2018). Studies using genotyping data of 84 SNPs indicated that the MoT isolates closely related to B71 were responsible for the wheat blast outbreaks in both Bangladesh and Zambia (Latorre and Burbano 2021; Win *et al*. 2021).

We previously generated the reference genome sequence for the B71 isolate, which has seven indispensable core-chromosomes and one supernumerary dispensable mini-chromosome (Peng *et al*. 2019). Supernumerary chromosomes, those which are not required for normal physiological growth and development, and which are often non-uniformly present in a population, exist in many plants, animals, and other fungi (Soyer *et al*. 2018; Bertazzoni *et al*. 2018; Yang *et al*. 2020). Supernumerary chromosomes are also called B-chromosomes, dispensable chromosomes, extra chromosomes, or accessory chromosomes. In *M. oryzae*, supernumerary chromosomes were typically smaller than the core chromosomes, consisting of a few megabases of DNA sequences, and they were therefore referred to as mini-chromosomes (Talbot *et al*. 1993; Orbach *et al*. 1996; Peng *et al*. 2019; Langner *et al*. 2021). The supernumerary chromosome is hypothesized to be an accelerator for fungal adaptive evolution (Coleman *et al*. 2009; Croll *et al*. 2013). Compared to core-chromosomes, supernumerary mini-chromosomes in *M. oryzae* have a lower gene density and higher levels of repetitive sequences. The repeat-rich mini-chromosome provides abundant intrachromosomal homology for DNA duplication, loss, and rearrangements, which can serve as a reservoir to accelerate fungal genome evolution (Peng *et al*. 2019). Indeed, sequence comparisons among mini-chromosomes from different *M. oryzae* strains showed that they are highly variable (Peng *et al*. 2019; Langner *et al*. 2021). For Bangladeshi and Zambian isolates that are genetically similar to B71, it is unknown if they carry mini-chromosomes and, if so, how similar they would be compared to the B71 mini-chromosome.

Strains in the wheat-adapted MoT lineage are still evolving and becoming more aggressive pathogens of wheat (Cruppe *et al*. 2020; Valent *et al*. 2021). The MoT population is still in the initial stages of moving from its center of origin into additional wheat regions of the world (Malaker *et al*. 2016; Islam *et al*. 2016; Tembo *et al*. 2020). Understanding and contrasting the evolutionary potential of MoT pathotype strains in the South American center of diversity and in the recently-emerged bottleneck populations in Bangladesh and Zambia, is a critical step to understand the movement and adaptation of wheat blast. In this study, we analyzed both single nucleotide polymorphisms (SNPs) and genomic structural variation among isolates from Bangladesh, Zambia, and South America. Further, Oxford Nanopore long-read sequencing and assembly were conducted for two Zambian isolates and used for genome comparisons among Zambian strains and B71. Our results show that all the analyzed isolates contained at least one mini-chromosome, and we document the high-speed creation of genetic variability within the mini-chromosome from closely related isolates.

## RESULTS

### Establishment of a finished reference genome of South American isolate B71

The previous reference B71 genome (B71Ref1), assembled from PacBio long reads, contained 22 gaps in core-chromosomes and five scaffolds from the mini-chromosome (Peng *et al*. 2019). To generate a complete genome assembly, we produced >250x Oxford Nanopore long reads with an N50 of 29 kb and the longest read of 166 kb (**Figure S1)**. The resulting assembly, produced using the Canu assembler and polishing with both Nanopore reads and Illumina reads, resulted in telomere-to-telomere assemblies of chromosomes 2 to 7, chromosome 1 with telomere repeat sequence on one end, a telomere-to-telomere mini-chromosome, and a circularized mitochondrial genome (**Figure 1, Table S1**) (Walker *et al*. 2014; Koren *et al*. 2017). Telomeric retrotransposons MoTeRs with various copy numbers were found in all telomeric regions, including on the mini-chromosomes (Starnes *et al*. 2012). Notably, one MoTeR insertion was identified approximately 759 kb away from the end of chromosome 3. The unfinished assembly at the beginning of chromosome 1 is presumably due to the high copy number of ribosomal DNA repeats (rDNA). The accuracy of assembly sequences is estimated to be >99.99% (**Table S2**) (He *et al*. 2020). The length of the finished mini-chromosome is 1.9 Mb, consisting of 51.7% repeats. Core-chromosomes contain less repetitive sequences and different core-chromosomes vary in their repetitive sequence content. Chromosomes 2, 4, and 5 contain repeats ranging from 3.7% to 4.5%, while chromosomes 1, 3, 6, and 7 contain higher levels of repeat content, ranging from 9.8% to 17.9% (**Table S1**). Centromere sequences can be inferred from all chromosomes, including the mini-chromosome (**Figure 1**) (Yadav *et al*. 2019). Genome annotation with the updated reference genome results in 11,864 protein coding genes, which contain homologs with 25 known effector genes (**Supplementary Data 1**).

**Figure 1.**
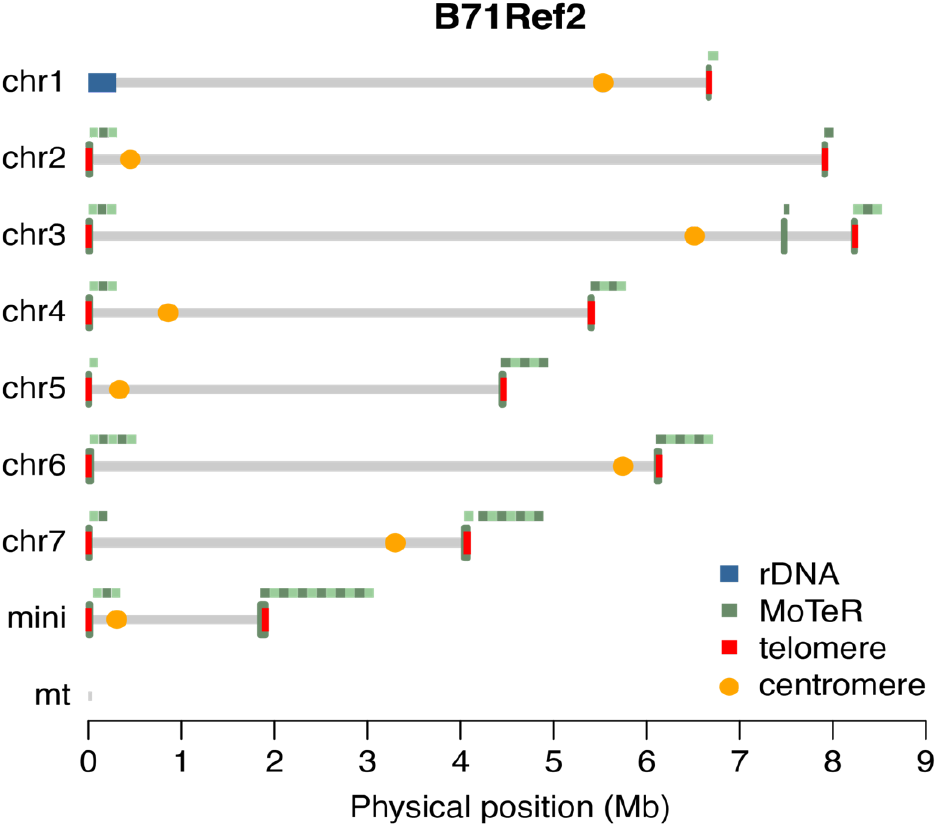
Chromosomal ends and centromeres of B71Ref2. The finished B71 reference genome (B71Ref2) consists of seven core-chromosomes, one supernumerary mini-chromosome, and a circularized mitochondrial genome. Positions of ribosomal DNA (rDNA) repeats, transposable element MoTeR repeats, telomere featured sequences, and inferred centromeres are highlighted. The close-up of each MoTeR repeat locus is shown above the locus with two green colors to indicate the copy number of MoTeR repeats.

### The B71 branch contains isolates from Bangladesh, Zambia, Bolivia, and Brazil

We collected publicly available whole genome sequencing (WGS) data of MoT strains, including Bangladeshi isolates collected from 2016 to 2020, Zambian isolates from 2018 to 2020, and isolates from South America. Phylogenetic analysis confirmed that all Bangladeshi and Zambian isolates cluster closely with B71 (**Figure 2a**) (Latorre and Burbano 2021; Win *et al*. 2021). We also identified a strain, 12.1.181, collected in 2012 from Brazil (Castroagudín *et al*. 2016), in the same phylogenetic clade as B71 and isolates from Bangladesh and Zambia (**Figure 2a**). Here, the clade is referred to as the B71 branch. Within the clade, Bangladeshi and Zambian isolates were separately clustered (**Figure 2b**). The Brazilian strain 12.1.181 is more similar to Bangladeshi isolates and the Bolivian strain B71 is more similar to Zambian isolates. These results indicate that the wheat blast outbreaks on the Asian and African continents were likely caused by different sub-branches of isolates. We identified 41 SNPs at which all Bangladeshi isolates shared one genotype, and 12.1.181 and B71 shared another (**Supplementary Data 2**). For all Zambian strains, none of these SNP sites harbored the Bangladesh isolate genotype, indicating that Zambian isolates spread from South America rather than from Bangladesh.

**Figure 2.**
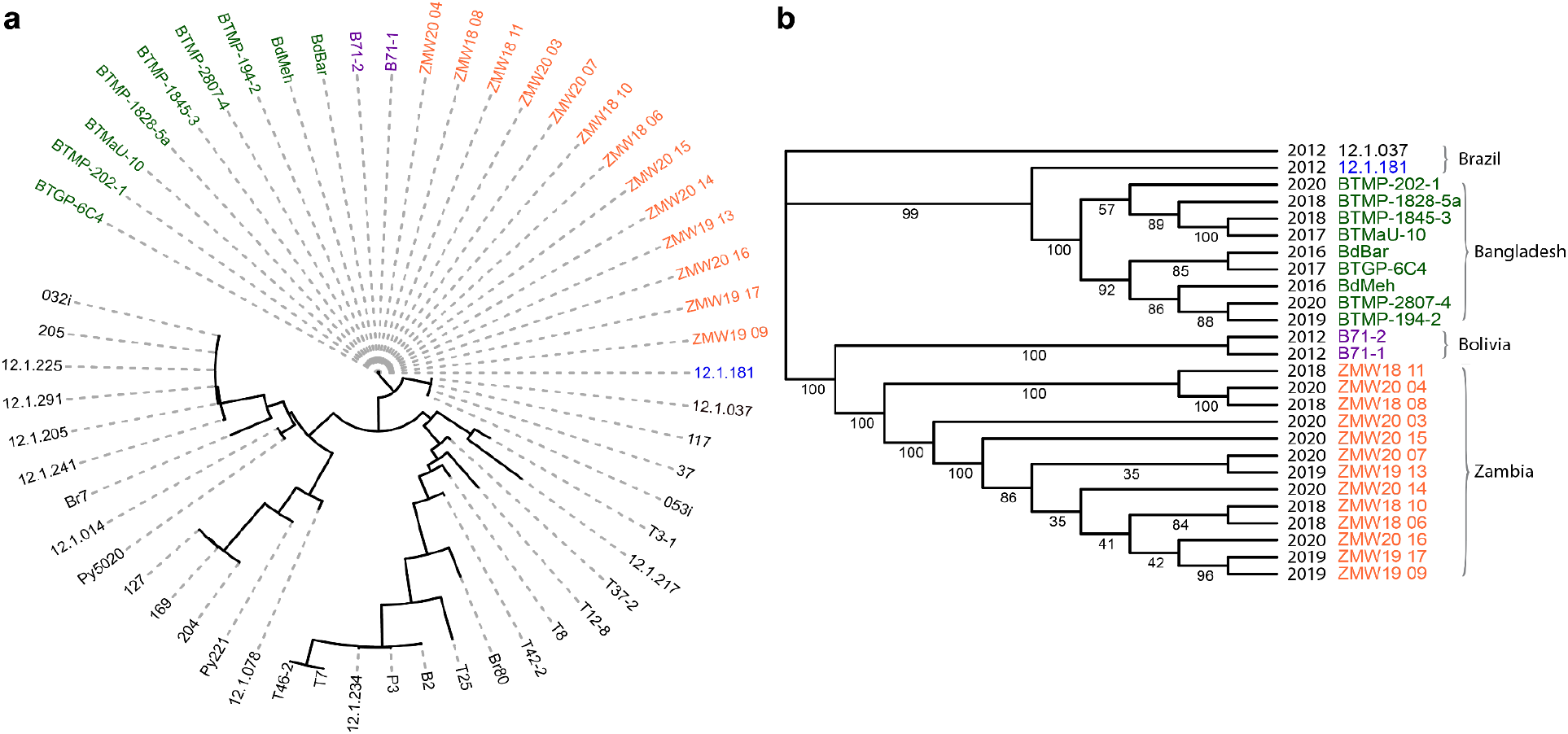
Phylogenetic tree of MoT isolates. Publicly available whole genome sequencing data of MoT isolates were used to identify SNPs for the phylogeny construction using the maximum likelihood approach. (**a**) Scaled tree of MoT isolates. B71-1 and B71-2 represent two independent samples of B71. (**b**) Unscaled tree of isolates from the B71 branch. Brazil isolate 12.1.037 served as the outgroup isolate. Numbers on edges are Bootstrap values. The year and country isolated for each isolate are labeled.

### Genomic structural variation among B71-related isolates

We performed Comparative Genomics Read Depth (CGRD) analysis against the new B71 assembly to estimate copy number variation using WGS read depth data (Lin *et al*. 2021). Given the relatively low sequence divergence between the isolates, as measured by SNPs, we were interested in assessing if copy number variation, including presence/absence variation and duplications, had taken place among the B71-related isolates. A few sequence segments that were present in B71 but absent in some other isolates were observed. This included a 155.5 kb segment at the end of chromosome 3 and a 22.5 kb segment at the beginning of chromosome 4 that were both absent in all Bangladeshi isolates and the Brazilian isolate 12.1.181 (**Figure 3a, 3b**). Both segments were present in all Zambian isolates. The results indicate the identified presence/absence variants were standing variation in South America prior to the Bangladesh introduction. The results also support the hypothesis that Zambian isolates are from a different sub-branch than those causing the Bangladesh outbreak.

**Figure 3.**
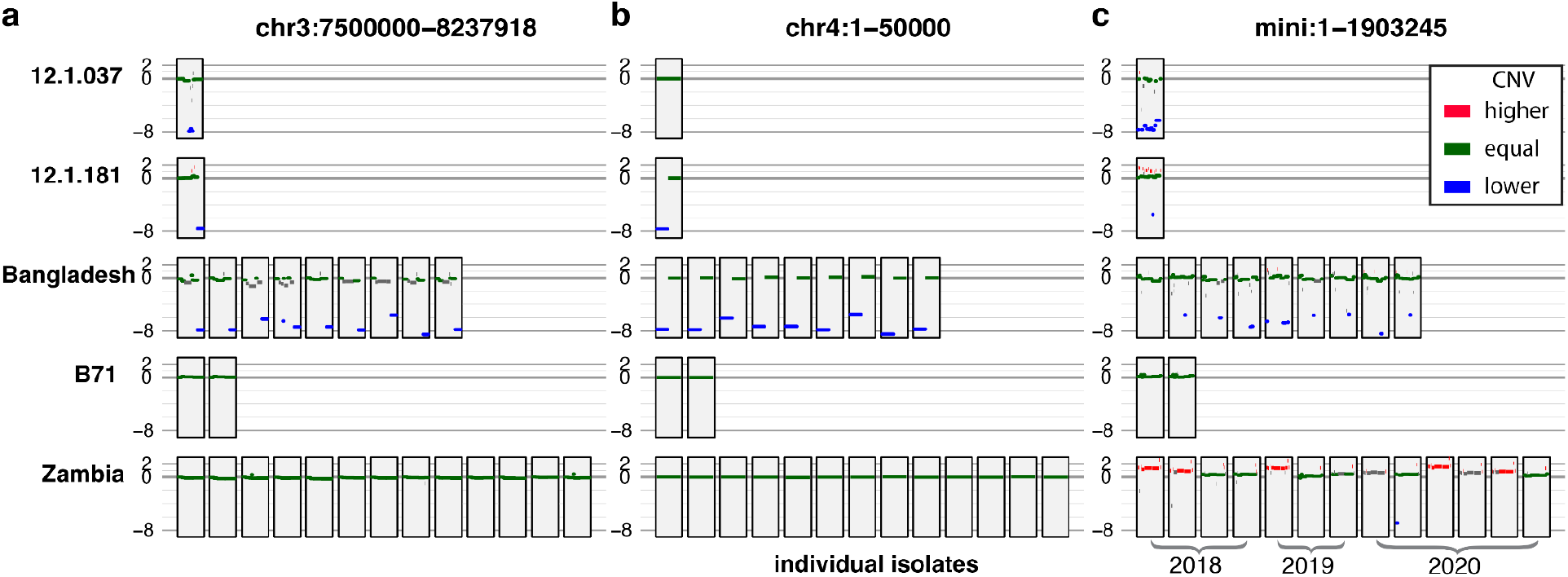
Regional results of CGRD. Each block represents the CGRD result at a region indicated on the top of each plot for an *M. oryzae* isolate. The CGRD result in each block was from the comparison between whole genome sequencing data of an isolate and B71. Results from two small regions on chromosome 3 (**a**) and chromosome 4 (**b**), the whole mini-chromosome (**c**) were displayed. The Y-axis values represent log2 values of ratios of read depths of the isolate to B71, signifying copy number variation (CNV). A value higher than 0.8, colored as red, indicates a higher copy number in the isolate as compared to B71. A value smaller than −5, colored as blue, indicates the absence of the B71 sequence in the isolate. The value between −0.4 and 0.4, colored as green, is deemed to be an equal copy number between the two genomes. Other values are colored as gray to indicate ungrouped regions. Years isolated for Zambian strains are labeled in (**c**).

In contrast to core-chromosomes, pervasive structural variation on the mini-chromosome was observed among isolates in the B71 branch (**Figure 3c**). All analyzed isolates from the branch contained most sequences of the B71 mini-chromosome, indicating that all these isolates carry a mini-chromosome similar to the originally described B71. As a control, we mapped reads from isolate 12.1.037, which was collected from Brazil, but is more genetically distinct from B71. As expected, we found that 12.1.037 has little sequence homology to the B71 mini-chromosome (**Figure 3c**). A few sequence regions of the B71 mini-chromosome were absent in the Bangladeshi isolates and 12.1.181 (**Figure 3c**). Two regions of the B71 mini-chromosome, approximately 6 kb and 86 kb at mini-chromosome positions 1.173 to 1.179 Mb and 1.364 to 1.450 Mb, respectively, were repeatedly found to be absent in genomes of multiple Bangladeshi isolates. The former region is also absent in the 12.1.181 genome. Only one region, from 101 kb to 588 kb, was absent in one of 13 Zambian isolates. CGRD analysis indicates that all Bangladeshi isolates carry one mini-chromosome, while Zambian isolates carry one or multiple mini-chromosomes (**Figure 3c**). Greater than one mini-chromosome was observed in Zambian strains isolated from Mpika in 2018, from Mount Makulu in 2019, and from multiple areas in 2020 (**Figure 3c**). The isolates containing either a single mini-chromosome or multiple mini-chromosomes were not clustered separately in the phylogenetic tree using whole genome SNP markers (**Figure 2b**).

### Rapid sequence divergence in the mini-chromosome of Zambian isolates

To confirm the observed mini-chromosome copy number dynamics inferred from short-read sequences of Zambian isolates, we conducted Oxford Nanopore long-read sequencing of two Zambian isolates, ZM1-2 and ZM2-1. The isolates were collected from the Mpika district of Muchinga Province, Zambia in 2018, which were previously termed as Zambia1.2 (ZM1-2) and Zambia2.1 (ZM2-1) (Tembo *et al*. 2020). The assembly resulted in nearly finished assemblies on all core-chromosomes for the two isolates (**Table S3, Figures S2, S3**). Both ZM1-2 and ZM2-1 assemblies consisted of ∼45 Mb with seven core-chromosomes and mini-chromosomal sequences. ZM2-1 has one telomere-to-telomere mini-chromosome sequence, while ZM1-2 has one telomere-to-telomere mini-chromosome sequence (termed mini1) and the other one with telomere-featured repeats on one end (termed mini2) (**Table S3, Figures S2, S3**).

Core-chromosomes of the two genomes are highly similar to the B71 core-chromosomes with a few large structural variants. Three hundred-kilobase insertion or deletion events in either ZM1-2 or ZM2-1 were identified (**Figure 4a, Figure S4, Table S4**). In addition, both chromosomes 6 and 7 have multiple smaller presence/absence variants among these three strains. Besides large insertions and deletions, inversions were identified on core-chromosomes. An inversion with more than 65 kb was found at around 3.3 Mb on chromosome 6 in both Zambian strains as compared to B71. A smaller inversion (>11 kb) was identified at around 0.39 Mb in ZM1-2 as compared with both ZM2-1 and B71 (**Table S5**). Overall, we estimated that the identified structural variants impacted less than 2% of core-chromosomes and largely occurred in gene-poor regions, possibly mediated by repetitive sequences.

**Figure 4.**
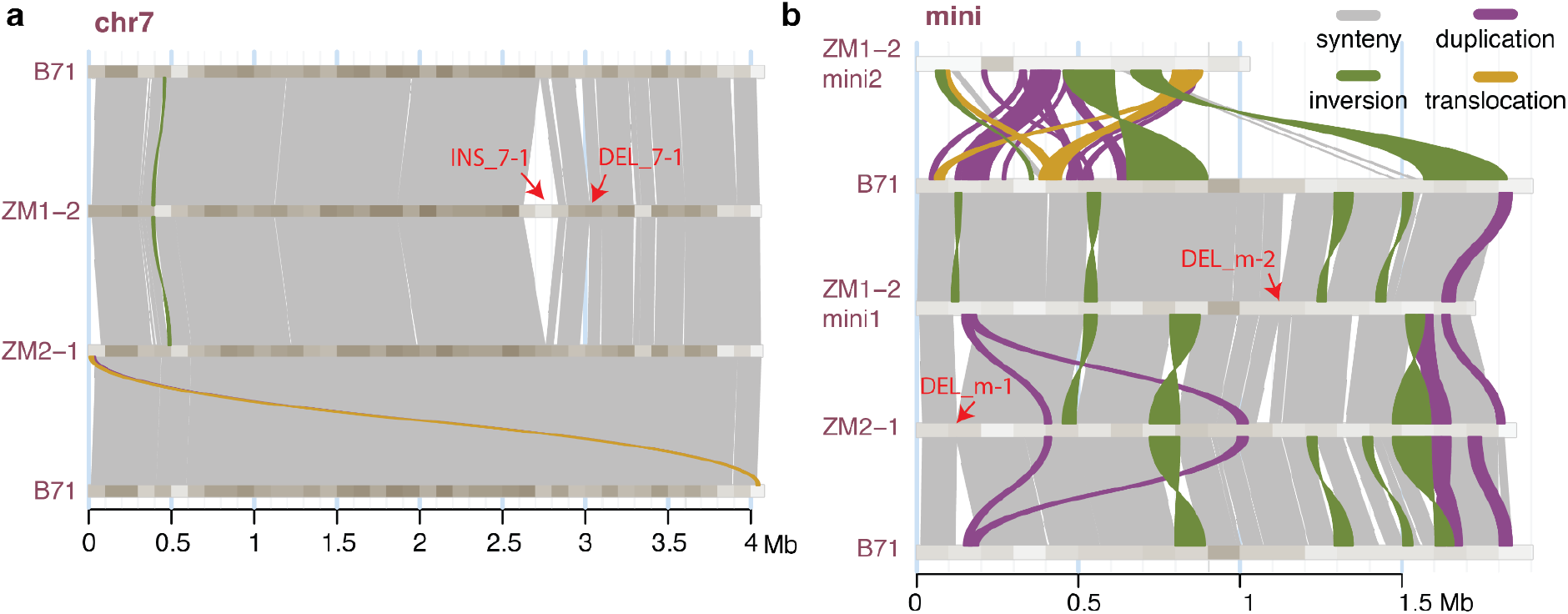
Comparison of chromosome 7 and mini-chromosomes among strains. (**a, b**) Chromosomal comparisons based on Syri analysis of structural variation. Alignments of synteny duplication, inversion, and translocation between chromosomes are color-coded. The gradient colors from tan to gray on chromosomes signify gene density from high to low. ZM1-2m1 and ZM1-2m2 are mini-chromosomes 1 (mini1) and 2 (mini2) of ZM1-2, respectively. Red errors point at large insertion (INS) and deletion (DEL) events. See Figure S4 for comparisons of chromosomes 1-6.

Consistent with our previous analysis using short reads of Zambian strains, long-read genome assemblies of both ZM1-2 and ZM2-1 have one mini-chromosome that is similar to the B71 mini-chromosome. The B71-like mini-chromosomes from Zambian strains contained multiple large presence/absence variants, inversions, duplications, and translocations (**Figure 4b**). For example, six inversion events were identified, indicative of frequent inversion events occurring within the B71 branch (**Figure 4b, Table S5**). Although large structural variation exists among B71 or B71-like mini-chromosomes, all of them carry the two effector genes *BAS1* and *PWL2* in mutually syntenic regions (**Figure 5**). Note that, using whole genome sequencing data of isolates with sufficient sequencing depths, reads with a nearly full coverage on both *BAS1* and *PWL2* were identified from all examined isolates of the B71 branch (**Figure S5**).

**Figure 5.**
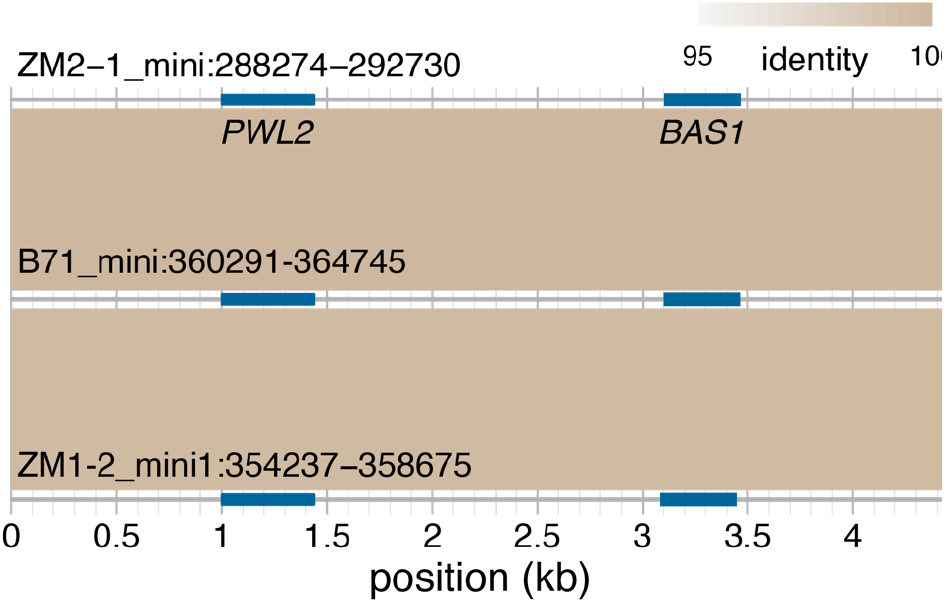
Syntenic alignments of the regions containing *PWL2* and *BAS1*. Both effector genes *PWL2* and *BAS1* are present in B71, ZM1-2 and ZM2-1. The regions harboring these two genes are nearly identical. Blue color bars signify the locations of the two genes and the bisque color indicates alignments between regions whose coordinates on the mini-chromosome are listed.

Interestingly, we also assembled an additional mini-chromosome (mini2, 1.03 Mb in size) from ZM1-2 that is distinct from the B71 mini-chromosome. As compared with the B71 mini-chromosome, ZM1-2 mini2 showed little chromosomal collinearity (**Figure 4b**). No known effector genes, including *BAS1* and *PWL2*, were found on mini2. The composition of transposable elements in ZM1-2 mini2 is not very different from that of other mini-chromosomes (**Figure S6**). All mini-chromosomes carry a relatively high proportion of retrotransposons, particularly, *Gypsy* LTR elements. ZM1-2 mini2 appears to contain less *TC1_Mariner* and *copia*, but more *helitron* as compared with other mini-chromosomes (**Figures S2, S6**).

## DISCUSSION

*M. oryzae* isolates responsible for the outbreaks of wheat blast in Bangladesh and Zambia were found to be related to a highly aggressive wheat blast strain B71 from South America (Latorre and Burbano 2021; Win *et al*. 2021). Here, we examined genomic variation among isolates of the B71 branch using an improved B71 reference genome. We constructed phylogenetic trees using SNPs and performed structural variation analyses using whole genome sequencing Illumina reads and genome assemblies with Nanopore long reads. Our results support that different sub-branches from South America were independently introduced to Bangladesh and Zambia. Genome sequencing data indicated that at least one mini-chromosome was maintained in all the analyzed isolates, providing the opportunity for examination of genomic variation of both core- and mini-chromosomes that diversified in a short evolutionary period.

### Isolates of different sub-branches caused outbreaks in Bangladesh and Zambia

Phylogenetic analysis using SNP genotyping data showed that Bangladeshi and Zambian isolates are closely related to MoT isolates B71 from Bolivia and 12.1.181 from Brazil. The phylogeny showed that all Bangladeshi isolates are closer to 12.1.181 and all Zambian isolates are closer to B71, indicative of separate sub-branches causing the outbreaks in Bangladesh and Zambia. We identified the 41 SNPs with a genotype conserved in all Bangladeshi strains, which were isolated from multiple cities in multiple years, and consistently different in all Zambian strains examined. The SNPs were located on all chromosomes, providing evidence that the introduction of wheat blast into Zambia occurred from South America rather than from Bangladesh. This finding is consistent with the conclusions from a recent independent analysis (Latorre *et al*. 2022). The hypothesis of independent wheat blast introductions to the two continents from South America was further supported by presence/absence polymorphisms on chromosomes 3 and 4. Importantly, for these two loci, both presence and absence genotypes existed in strains from South America, implying that the diversification of the ancestors of Bangladeshi and Zambian isolates occurred in South America. In conclusion, our genomic analyses strongly support that isolates from different sub-branches from South America caused the first outbreaks in Bangladesh and Zambia.

### The mini-chromosome has been maintained, but is rapidly diversifying

Our results indicate that all field isolates from the B71 branch carry at least one mini-chromosome, and some Zambian strains carry more than one. The maintenance of the mini-chromosome in all examined isolates indicates that depletion of the mini-chromosome rarely occurred and/or selection pressure existed for retaining the mini-chromosome. The mini-chromosome harbors more than 50% repetitive sequences, including a high proportion of retrotransposons and DNA transposons, while core-chromosomes contain ∼10% repetitive sequences and a higher density of genes. What selection could be acted upon to maintain these otherwise dispensable sequences remains to be determined. We observed pervasive large structural variation among mini-chromosomes from different isolates in the same branch, in contrast with the relatively infrequent structural changes found between core-chromosomes. Although there is pervasive structural variation, some genes in mini-chromosomes appear to be well retained in all strains. Whole genome sequencing data showed that all examined strains of the B71 branch contain sequences of the effector genes *BAS1* and *PWL2* (Sweigard *et al*. 1995; Mosquera *et al*. 2009), indicating the genomic region containing the two genes is relatively stable despite so far only being localized to highly dynamic mini-chromosomes in the MoT population. Additionally, *BAS1* and *PWL2* co-occur in 6% of wheat isolates collected between 1986 and 1989, and in 91% of wheat isolates collected in 2017 and 2018, indicating that the overall frequency of wheat field isolates containing these two genes has increased through time (Navia-Urrutia *et al*. 2022). These results suggest that *BAS1* and *PWL2* might be playing a role in the enhanced aggressiveness of more recent MoT strains towards wheat. The presence of *BAS1* and *PWL2* in all examined strains also implies that these strains likely contain the B71-like mini-chromosome because the mini-chromosome (mini2) that is highly divergent from the B71-like mini-chromosome did not carry these two genes.

Multiple mini-chromosomes were found in individual Zambian strains isolated in multiple years (2018, 2019, and 2020). Our analysis did not provide evidence that Zambian strains with more than one mini-chromosome have increased in frequency, but this result is limited due to the limited number of Zambian strains collected to date. The additional mini-chromosome (mini2) we found in Zambian isolate ZM1-2 contained no known effector genes. We speculate that the high variability of the observed mini2 is not necessarily representative of all mini-chromosome variation, and there may be additional significant mini-chromosome variation in terms of copy number and sequence content. It is intriguing to observe mini-chromosome variation occurring in a natural population in a matter of years. While copy number variation is a well-documented means to increase biotic and abiotic stress tolerance (Cook *et al*. 2012; Maron *et al*. 2013), most examples are linked to copy number changes in the core genome. To our knowledge, there are few examples of rapid sequence divergence of non-core genomic compartments in natural populations. Recent analysis of naturally occurring herbicide resistant *Amaranthus palmeri*, an economically important weed impacting crop production, found significant expansion of the coding sequence for 5-enolpyruvylshikimate-3-phosphate synthase (*EPSPS*) occurring on extrachromosomal circular DNA (eccDNA) (Koo *et al*. 2018). The importance of accessory chromosomes is clear in plant pathogenic fungi, where they can be transferred and affect the host-range and/or habitat diversity of individual strains of *Fusarium* spp. (Coleman *et al*. 2009; Ma *et al*. 2010). The biological impact of the documented mini-chromosome divergence described here remains to be determined. Likewise, it is not clear if there is a genetic or environmental factor contributing to the mini-chromosome divergence found in strains collected in Zambia, or if this is a matter of ascertainment bias and represents a common phenomenon in *M. oryzae*. Analysis of additional strains from the branch in coming years will help further address why the B71-like mini-chromosome was better maintained in the field than the other one (mini2), whether the mini-chromosome can be depleted in field strains, and how multiple mini-chromosomes in one strain arose.

## METHODS

### Genome assembly of B71

Nanopore FASTQ read data were generated using basecaller Guppy (version 3.4.4) with default parameters (http://nanoporetech.com/community). Reads were assembled with Canu (version 1.9) with parameters (genomeSize=45m minReadLength=5000 minOverlapLength=1000 corOutCoverage=80). The previous assembly version (B71Ref1.6) was used to add missing chromosome ends (He *et al*. 2020). A contig containing ribosomal DNA sequences was added to the beginning of chromosome 1. The resulting assembly was polished using Nanopolish with Nanopore raw FAST5 data and Pilon with Illumina reads (SRA accession: SRR6232156) (Walker *et al*. 2014; Loman *et al*. 2015).

### Public Illumina sequencing data

Publicly available data of MoT strains were downloaded from Sequence Read Archive (SRA). Accession numbers are available in **Table S6**.

### Illumina whole genome sequencing (WGS)

All MoT strains are maintained in frozen storage and manipulated under Biosafety Level 3 (BSL3) laboratory conditions in the USDA-ARS Foreign Disease and Weed Science Research Unit (FDWSRU) in Fort Detrick, Maryland and in the Biosecurity Research Institute (BRI) at Kansas State University in Manhattan, KS, as authorized by PPQ permits from the USDA-Animal and Plant Health Inspection Services. *Magnaporthe* isolates were cultured on oatmeal agar (OMA) plates seeded with dried Whatman filter paper containing mycelia and conidia (Valent *et al*. 1986). The culture plates were incubated under continuous light at room temperature for 7-12 days. Liquid cultures were seeded with 1 cm^2^ piece of OMA culture from the actively growing region of the plate into minimal medium (3 g of Casamino acids, 3 g of yeast extract, and 6 g of sucrose in 1 liter of H2O) at 24°C in a stationary flask. After seven days, mycelial mats from each sample were collected, blotted dry on paper towels, lyophilized for 24 hr and stored at room temperature. DNA extraction was performed by finely grinding 200 mg of lyophilized mycelium in liquid nitrogen. Five hundred microliters of extraction buffer (1% CTAB, 0.7 M NaCl, 100mM Tris [pH 7.5], 10 mM EDTA, 1% 2-Mercaptoethanol, 0.3 mg/ml Proteinase K) was added and mixed thoroughly by shaking and tube inversion. The mixture was incubated at 65°C for 30 mins, then allowed to cool to room temperature. One milliliter of phenol:chloroform:isoamyl alcohol (25:24:1) was added and mixed by shaking and inverting tubes. The tubes were centrifuged for 10 mins at 14,000 rpm in a benchtop centrifuge. The aqueous phase was removed, and the DNA was precipitated by mixing in 0.54 volumes of room temperature isopropanol, followed by centrifugation at 14,000 rpm for 10 minutes. The pellets were rinsed with 70% ethanol and then allowed to air dry. Pellets were dissolved in 50 ml of TE buffer containing 1 mg/ml RNase. Paired-end sequencing was performed on the isolates using the Illumina platform at Novogene USA.

### Nanopore WGS for de novo genome assemblies of ZM1-2 and ZM2-1

Genomic DNA was extracted from the mycelial powder using a modified CTAB method (He 2000). To filter small-size DNA fragments, BluePippin Gel Cassette (Sage Science, USA, Cat.# BLF7510) was used to perform a >20 kb High-Pass size selection. About 1 ug of the selected DNA was used to build a library using SQK-LSK110 sequencing kit (Oxford Nanopore, UK, Cat.# SQK-LSK110). Twelve microliters of the DNA library was loaded to a R9.4.1 flow cell (Oxford Nanopore, UK, Cat.# FLO-106D) and performed whole genome sequencing on a MinION Sequence Device (Oxford Nanopore, UK, Cat.# MinION Mk1B). The Nanopore raw data stored in FAST5 format was converted to FASTQ using Guppy Basecaller (version 6.0.1, https://community.nanoporetech.com). Genome was assembled using Canu (version 2.2, https://github.com/marbl/canu) (Koren *et al*. 2017) with the parameters of “minReadLength=5000 minOverlapLength=1500 corOutCoverage=80 correctedErrorRate=0.1”. The contigs in Canu assemblies were aligned to B71Ref2 to determine the chromosome number and the orientation using NUCmer (version 4.0.0, https://mummer4.github.io/) with the parameters of “-L10000 -I 90” (Marçais *et al*. 2018). For the strain ZM1-2, the contigs assembled by Canu were further assembled into two mini-chromosomes based on the Bandage visualization (Version 0.9.0, https://github.com/rrwick/Bandage) (Wick *et al*. 2015) with the Canu output (best.edges.gfa) and the Nucmer alignment among the 8 contigs. The resulting assembly was polished using Nanopolish (version 0.13.3) for two runs and Pillon (version 1.24) for two runs (Walker *et al*. 2014; Loman *et al*. 2015). For the strain ZM2-1 Nanopore sequencing, we noticed the contamination of *Neurospora crassa* based on the result from data analysis. Therefore, contigs matching the *Neurospora crassa* reference genome (NC_026501.1) were removed after Nanopolishing.

### Estimation of base errors in genome assemblies

Base errors were estimated based on software KAD (version 0.1.7) with the parameter of “--minc 5 --klen 35” (He *et al*. 2020).

### Annotation of transposable elements

Extensive *de novo* TE Annotator (EDTA, v2.0.0) was used for transposable element annotation (Ou *et al*. 2019).

### Genome annotation

Pipeline funannotate (version 1.8.8) was used for genome annotation. RNA-seq data from both plate culture and *in planta* infection were used as expression evidence (SRA accessions: SRR9126640, SRR9127597 to SRR9127602). Protein data include MG8 protein annotation from rice isolate 70-15 (Dean *et al*. 2005), UniProtKB/Swiss-Prot protein database released in March 2021, an effector collection (https://raw.githubusercontent.com/liu3zhenlab/collected_data/master/Magnaporthe/known.effectors.db01.fasta).

### SNP identification

Paired-end Illumina sequencing reads were trimmed with Trimmomatic (version 0.38) (Bolger *et al*. 2014). The resulting paired-end reads were aligned to the reference (B71Ref2) with BWA (version 0.7.17-r1188) (Li and Durbin 2010). Alignments were filtered to retain confident alignments with at least 60 bp matches, >95% identity, and >95% coverage per read. GATK (version 4.1.0.0) was used for SNP discovery and filtering (McKenna *et al*. 2010). We only remained biallelic SNP sites with the B71 isolate matching the B71 reference genome.

### Construction of phylogenetic trees

Phylogenetic trees were constructed using IQ-TREE (version 1.6.12) (Minh *et al*. 2020). Briefly, two steps were performed. The first step identified the optimized model, which was then used in the second step for tree construction.

### Analysis of copy number variation via CGRD

Pipeline CGRD (version 0.3.5) was used to find large copy number variation (e.g., presence/absence variation and duplication) using Illumina data with the adjusted parameters (--knum --adj0 --cleanup --groupval “-5 −0.4 0.4 0.8”) (Lin *et al*. 2021).

### Analysis of structural variation via Syri

Genome sequences of B71, ZM1-2, and ZM2-1 were aligned with NUCmer (version 4.0.0) with the parameters (--maxmatch -c 500 -b 500 -l 20) (Marçais *et al*. 2018). Alignments were filtered with “delta-filter” in NUCmer with the parameters (-m -i 90 -l 500). Filtered alignments were then input to SyRI (version 1.5) for structural variation analysis (Goel *et al*. 2019). Two mini-chromosomes of ZM1-2 were separately combined with core-chromosomes for whole genome alignments and SYRI analysis. The in-house R module “mgplot” was employed for displaying chromosomal alignments and large structural variation.

### Read coverages of *PWL2* and *Bas1*

Strains that are in the B71 branch and have at least 8 million pairs of reads were used for examining read coverages of *PWL2* and *BAS1*. T25 that has no mini-chromosomes and did not contain either of these two genes was used as the control. Briefly, trimmed Illumina reads were aligned to each gene using BWA (version 0.7.17-r1188) (Li and Durbin 2010). Alignments with at least 60 match, 98% identity, and 98% coverage were retained and read coverage per basepair position was determined and plotted.

## ACKNOWLEDGEMENTS

We thank Dr. Mark Farman from Department of Plant Pathology at University of Kentucky for valuable comments. S. Ramachandran was supported in part by an appointment to the Foreign Disease and Weed Science Research Unit, United States Department of Agriculture, administered by ORAU through the U.S. Department of Energy Oak Ridge Institute for Science and Education and USDA-ARS. Funding was provided by the USDA NIFA award (no. 2018-67013-28511) to S. Liu, and USDA NIFA award (2021-67013-35724) to S. Liu, B. Valent, D. Cook, and D. Koo, the NSF award (no. 1741090) to S. Liu, and NSF award (2011500) to S. Liu, B. Valent, D. Cook, and D. Koo. This research was supported by USDA-ARS projects 8044-22000-046-00D and 8044-22000-051-00D. This is contribution no. 22-309-J from the Kansas Agricultural Experiment Station, Manhattan, Kansas.

## AUTHOR CONTRIBUTION

KP, BV, and SL conceptualized experiments; GL, SR and GC conducted experiments; GL, DC, SR, and SL analyzed data; SL, GL, SR, DC, KP, and BV wrote the manuscript; all authors reviewed and revised the manuscript.

## DATA AVAILABILITY

Genomic sequencing and assembly data of ZM1-2 and ZM2-1 have been deposited in the Sequence Read Archive (SRA) database under accessions PRJNA849498 and PRJNA849580, respectively.

## CODE AVAILABILITY

Related scripts are available at GitHub (https://github.com/PlantG3/B71branch).

