## Supplementary Information for "Rapid mini-chromosome divergence among fungal isolates causing wheat blast outbreaks in Bangladesh and Zambia"

Liu *et al*.

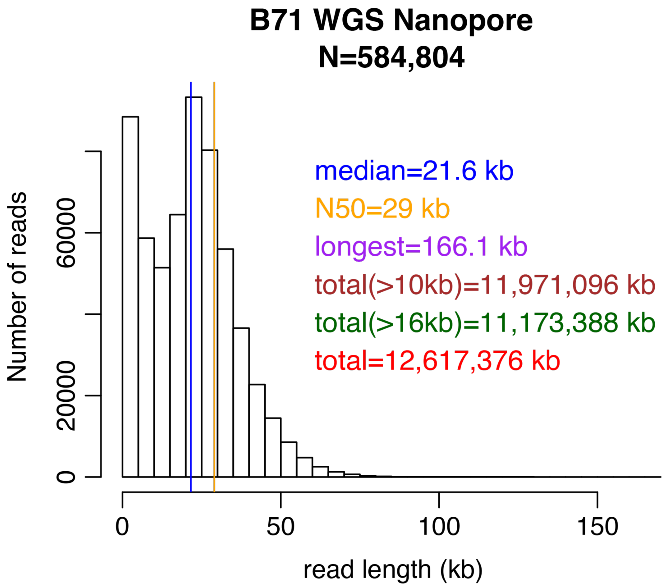

**Figure S1. Histogram of lengths of Nanopore raw reads**
Whole genome sequencing of B71 using a MinION flowcell produced ~12.7 Gb total sequences. The median and N50 of read lengths are indicated by blue and orange vertical lines, respectively.

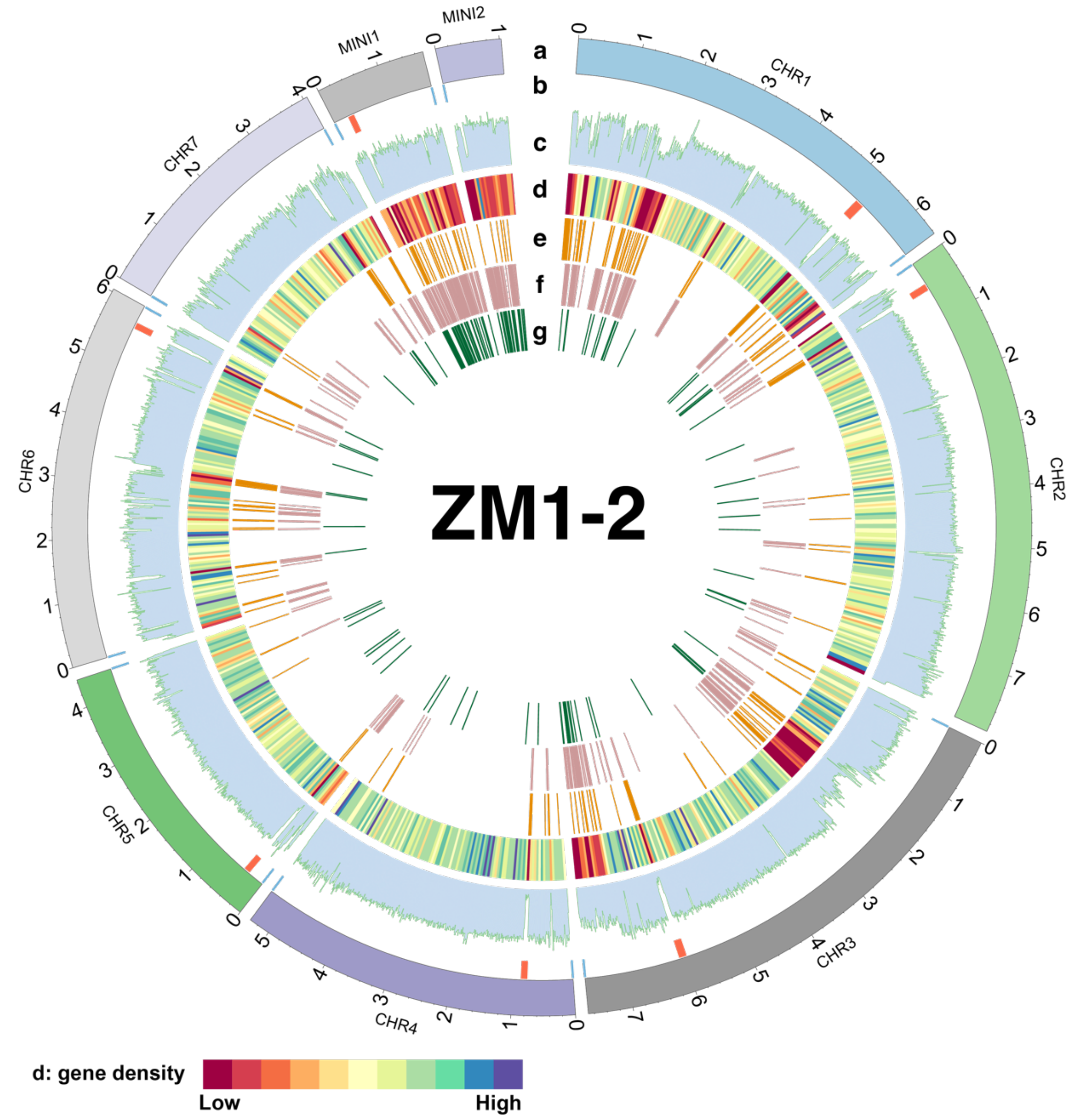

**Figure S2. Genomic features of ZM1-2**

(**a**) Core- and mini-chromosomes of ZM1-2. (**b**) Centromeres and telomeres on each chromosome. Blue lines indicate the locations of telomeric repeats (CCCTAA or TTAGGG), and red lines indicate the locations of centromeric regions matching the corresponding B71 centromeres with at least 10 kb and 90% identity. (**c**) GC content of the chromosomes. The average GC contents of 10 kb window sequences were calculated, and the minimum and maximum GC contents displayed on the histogram are 20% and 70%, respectively. (**d**) The gene density along chromosomes. The gene number per 50 kb window sequence is color-coded. (**e-g**) The locations of transposon elements of *Copia*, *Gypsy* and *CTCTA*. Each line on the track represents an annotated transposable element.

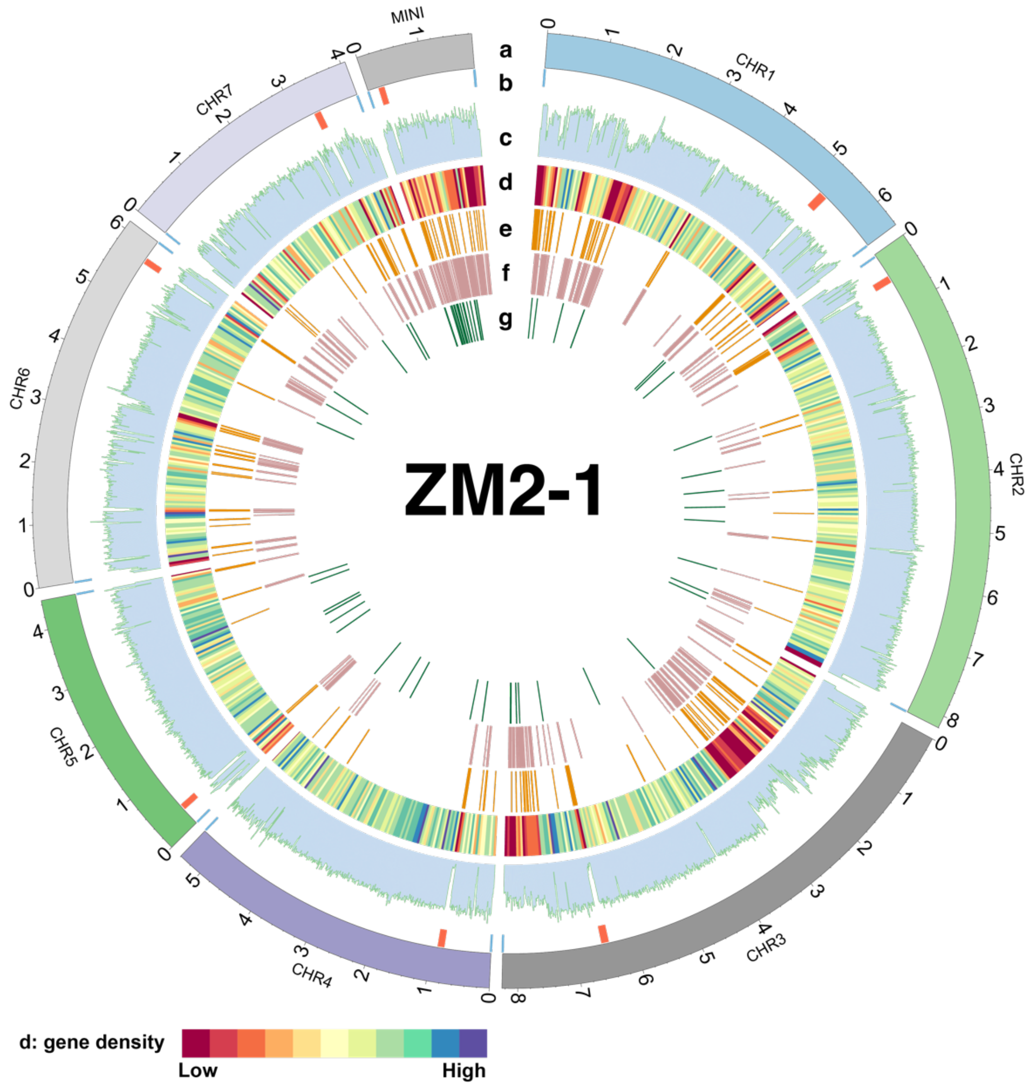

**Figure S3. Genomic features of ZM2-1**

(**a**) Core- and mini-chromosomes of ZM2-1. (**b**) Centromeres and telomeres on each chromosome. Blue lines indicate the locations of telomeric repeats (CCCTAA or TTAGGG), and red lines indicate the locations of centromeric regions matching the corresponding B71 centromeres with at least 10 kb and 90% identity. (**c**) GC content of the chromosomes. The average GC contents of 10 kb window sequences were calculated, and the minimum and maximum GC contents displayed on the histogram are 20% and 70%, respectively. (**d**) The gene density along chromosomes. The gene number per 50 kb window sequence is color-coded. (**e-g**) The locations of transposon elements of *Copia*, *Gypsy* and *CTCTA*. Each line on the track represents an annotated transposable element.

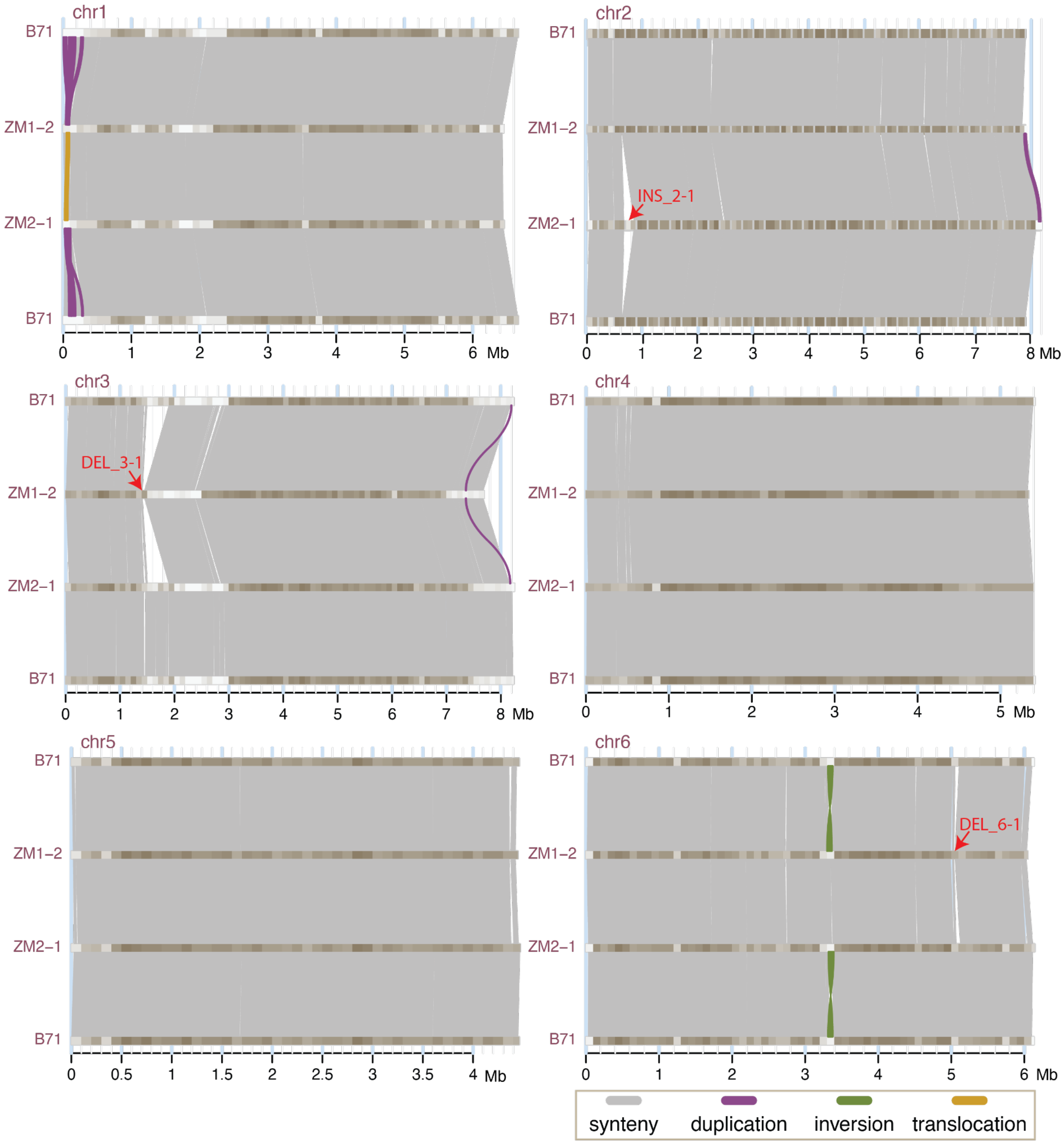

**Figure S4. Comparison of chromosomes 1-6 among strains**

Chromosomal comparisons based on Syri analysis of structural variation. Alignments of synteny duplication, inversion, and translocation between chromosomes are color-coded. Duplication and translocation at the beginning of chromosome 1 are likely artifacts due to poor assemblies of the highly repetitive ribosomal DNA region. The gradient colors from tan to gray on chromosomes signify gene density from high to low. Red errors point at large insertion (INS) and deletion (DEL) events.

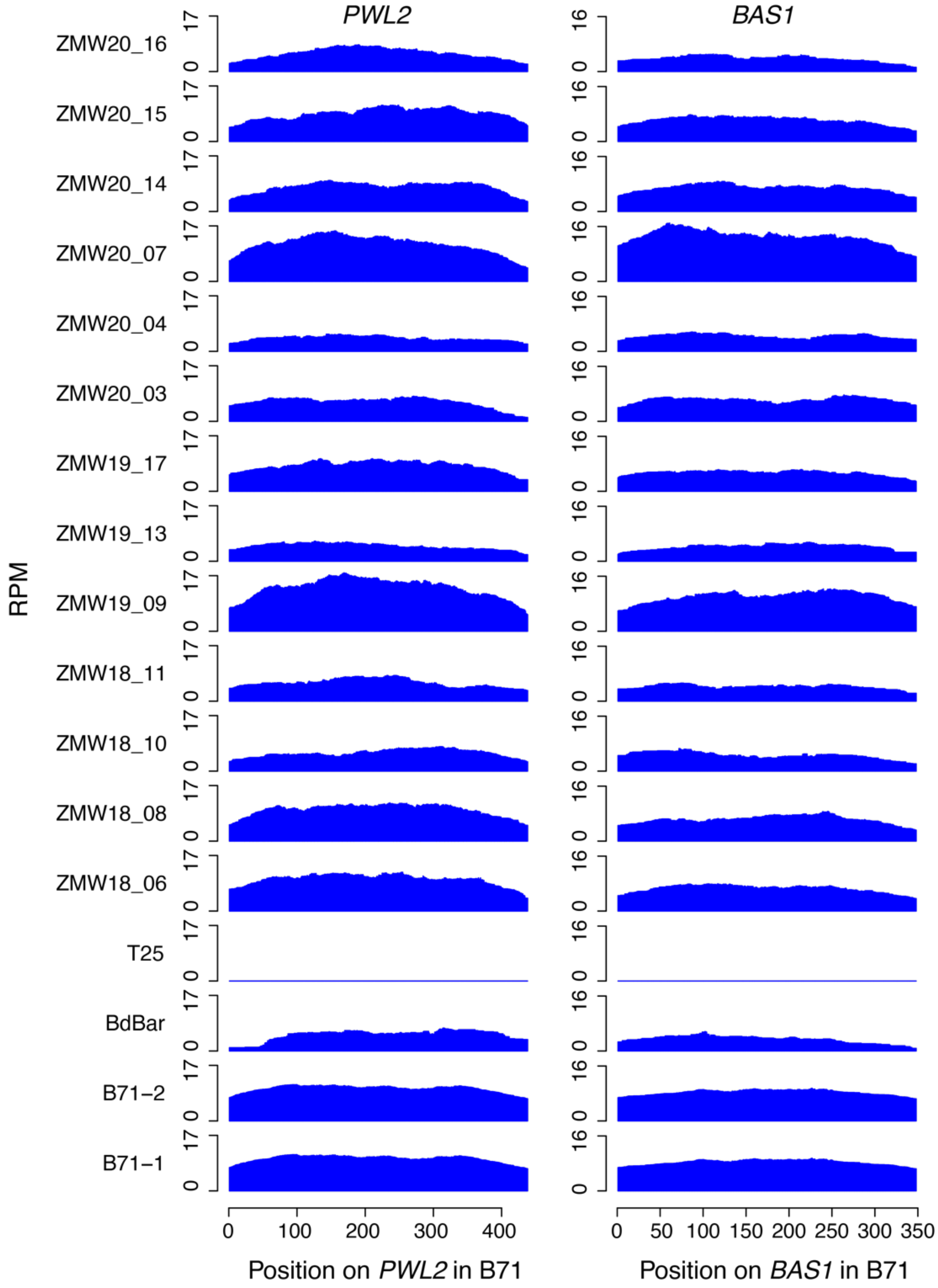

**Figure S5. Read coverages on *PWL2* and *BAS1***

Whole genome sequences of each isolate were aligned to *PWL2* and *BAS1* genes. Reads per million of total reads (RPM) were determined at each position and plotted. All isolates except T25 are from the B71 branch. T25 was previously verified to not carry either *PWL2* or *BAS1*, and no reads on either gene were identified. T25 has no mini-chromosomes.

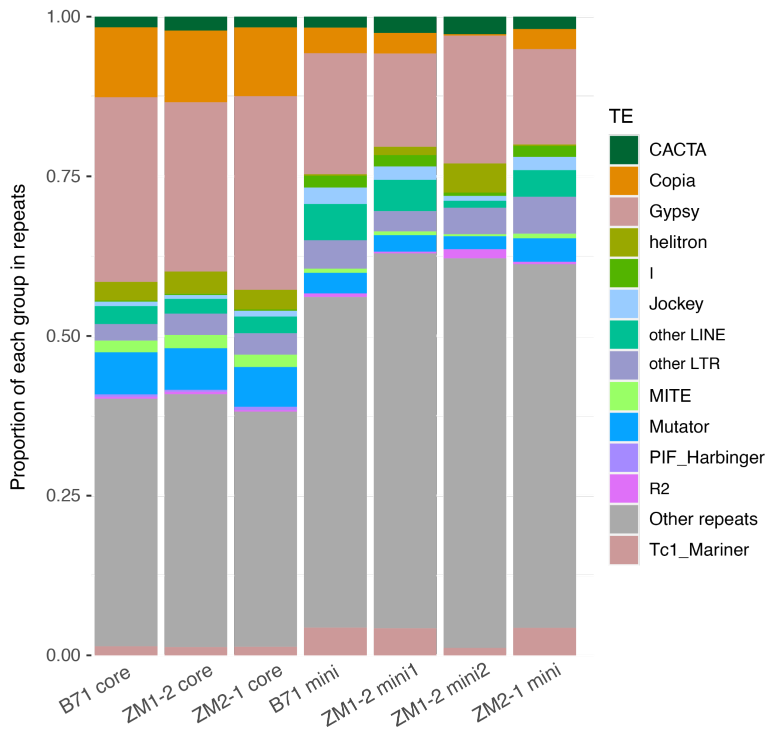

**Figure S6. Proportional distribution of transposon families by lengths**
The total length of each transposon family was determined for core-chromosomes (core) and mini-chromosomes (mini, mini1, or mini2) in each strain. For core-chromosomes and each mini-chromosome of a strain, the proportion of the total length of each transposon family out of the total transposon length from all families is displayed and stacked.

**Table S1.** Chromosome lengths and repeat contents of B71

| Chr | Length (bp) | # Repeats | Repeats (bp) | Repeats (%) | Telomere repeats* |
|---|---|---|---|---|---|
| chr1 | 6,668,242 | 1,204 | 1,194,042 | 17.9 | R |
| chr2 | 7,915,472 | 263 | 356,303 | 4.5 | L+R |
| chr3 | 8,237,918 | 1,337 | 1,287,291 | 15.6 | L+R |
| chr4 | 5,413,369 | 199 | 240,484 | 4.4 | L+R |
| chr5 | 4,460,276 | 150 | 165,579 | 3.7 | L+R |
| chr6 | 6,133,529 | 450 | 598,575 | 9.8 | L+R |
| chr7 | 4,079,358 | 369 | 460,817 | 11.3 | L+R |
| mini | 1,903,245 | 1,008 | 983,719 | 51.7 | L+R |
| mt | 34,996 | 1 | 115 | 0.3 | NA |

* L and R stand for telomeric repeats on the left and right sides, respectively. NA: not available

**Table S2.** Base accuracy of B71Ref2

| Assembly | Number_error_base* | Base_accuracy (%) |
|---|---|---|
| B71Ref1 | 7,553 | 99.98% |
| B71Ref1.6 | 3,057 | 99.99% |
| B71Ref2 | 2,164 | 100.00% |

* error bases estimated by KAD using k=35

**Table S3**. Chromosome lengths and repeat contents of ZM1-2 and ZM2-1

| Strain | Chr | Length (bp) | # Repeats | Repeats (bp) | Repeats (%) | Telomere repeats* |
|---|---|---|---|---|---|---|
| ZM1-2 | chr1 | 6,453,676 | 1,178 | 1,072,047 | 16.6% | R |
| ZM1-2 | chr2 | 7,922,507 | 266 | 315,652 | 4.0% | L |
| ZM1-2 | chr3 | 7,693,800 | 1,397 | 1,185,593 | 15.4% | L+R |
| ZM1-2 | chr4 | 5,338,394 | 180 | 176,663 | 3.3% | L+R |
| ZM1-2 | chr5 | 4,445,134 | 142 | 143,446 | 3.2% | L+R |
| ZM1-2 | chr6 | 6,045,076 | 466 | 524,838 | 8.7% | L+R |
| ZM1-2 | chr7 | 4,061,603 | 326 | 320,336 | 7.9% | L+R |
| ZM1-2 | mini1 | 1,726,145 | 1,222 | 1,032,342 | 59.8% | L+R |
| ZM1-2 | mini2 | 1,027,500 | 737 | 614,632 | 59.8% | L |
| ZM1-2 | mt | 34,946 | 1 | 115 | 0.3% | NA |
| ZM2-1 | chr1 | 6,459,530 | 1,061 | 1,017,943 | 15.8% | L+R |
| ZM2-1 | chr2 | 8,200,447 | 284 | 409,633 | 5.0% | L+R |
| ZM2-1 | chr3 | 8,249,511 | 1,342 | 1,260,143 | 15.3% | R |
| ZM2-1 | chr4 | 5,403,669 | 164 | 226,058 | 4.2% | L+R |
| ZM2-1 | chr5 | 4,468,408 | 132 | 171,465 | 3.8% | L+R |
| ZM2-1 | chr6 | 6,142,428 | 422 | 595,854 | 9.7% | L+R |
| ZM2-1 | chr7 | 4,070,502 | 342 | 446,126 | 11.0% | L+R |
| ZM2-1 | mini | 1,852,833 | 1,096 | 1,030,348 | 55.6% | L+R |
| ZM2-1 | mt | 35,002 | 1 | 115 | 0.3% | NA |

* L and R stand for telomeric repeats on the left and right sides, respectively. NA: not available

**Table S4**. Insertions or deletions with at least 40 kb

| SV | Strain | Length | Region* |
|---|---|---|---|
| INS_2-1 | ZM2-1 | 170,208 bp | ZM2-1_chr2:677153-847360 |
| INS_7-1 | ZM1-2 | 207,955 bp | ZM1-2_chr7:2628827-2836781 |
| DEL_3-1 | ZM1-2 | >340 kb | ZM2-1_chr3:1538919-1880542DEL; B71_chr3:1528859-1878215 |
| DEL_6-1 | ZM1-2 | >40 kb | B71-chr6:5055408-5104729; ZM2-1_chr6:5070036-5119358 |
| DEL_7-1 | ZM1-2 | >43 kb | B71_chr7:2936487-2979831; ZM2-1_chr7:2968888-3012233 |
| DEL_m-1 | ZM2-1 | >73 kb | ZM1-2_mini1:113509-186675; B71_mini:120194-192795 |
| DEL_m-2 | ZM1-2 | ~40 kb | B71_mini:1133643-1173428; ZM2-1_mini:1050305-1090093 |

* regions are expressed in the format with strain name, chromosome name, and two start and end coordinates.

**Table S5**. Large inversions among three strains

| Inversion | genome 1 | | | | genome 2 | | | | Unique* |
|---|---|---|---|---|---|---|---|---|---|
| | strain | chr | start (bp) | end (bp) | strain | chr | start (bp) | end (bp) | |
| INV_6-1 | B71 | chr6 | 3,312,253 | 3,377,306 | ZM1-2 | chr6 | 3,298,991 | 3,366,159 | B71 |
| INV_6-1 | B71 | chr6 | 3,312,253 | 3,377,306 | ZM2-1 | chr6 | 3,317,572 | 3,391,733 | B71 |
| INV_7-1 | B71 | chr7 | 452,947 | 464,499 | ZM1-2 | chr7 | 386,458 | 397,784 | ZM1-2 |
| INV_7-1 | ZM2-1 | chr7 | 482,938 | 494,490 | ZM1-2 | chr7 | 386,458 | 397,784 | ZM1-2 |
| INV_m-1 | ZM2-1 | mini | 720,192 | 811,320 | B71 | mini | 799,517 | 890,636 | ZM2-1 |
| INV_m-1 | ZM2-1 | mini | 720,192 | 811,320 | ZM1-2 | mini1 | 784,404 | 875,602 | ZM2-1 |
| INV_m-2 | ZM2-1 | mini | 1,472,127 | 1,589,223 | B71 | mini | 1,603,783 | 1,659,003 | ZM2-1 |
| INV_m-2 | ZM2-1 | mini | 1,472,127 | 1,589,223 | ZM1-2 | mini1 | 1,513,234 | 1,568,676 | ZM2-1 |
| INV_m-3 | B71 | mini | 1,292,262 | 1,348,447 | ZM2-1 | mini | 1,208,900 | 1,235,586 | B71 |
| INV_m-3 | B71 | mini | 1,292,262 | 1,348,447 | ZM1-2 | mini1 | 1,238,120 | 1,264,807 | B71 |
| INV_m-4 | B71 | mini | 1,504,842 | 1,531,822 | ZM2-1 | mini | 1,380,196 | 1,407,176 | B71 |
| INV_m-4 | B71 | mini | 1,504,842 | 1,531,822 | ZM1-2 | mini1 | 1,421,256 | 1,448,245 | B71 |
| INV_m-5 | ZM1-2 | mini1 | 517,855 | 556,237 | B71 | mini | 529,454 | 568,345 | ZM1-2 |
| INV_m-5 | ZM1-2 | mini1 | 517,855 | 556,237 | ZM2-1 | mini | 451,861 | 492,081 | ZM1-2 |
| INV_m-6 | ZM1-2 | mini1 | 109,242 | 128,692 | B71 | mini | 120,201 | 139,631 | NA[&] |

* the one from B71, ZM1-2, and ZM2-1 contains an inversion as compared with the other two

[&] the inversion only identified between mini1 of ZM1-2 and mini of B71 due to the region was absent in ZM2-1

**Table S6**. SRA accessions of publicly available WGS data

| Isolate | SRA_accession |
|---|---|
| 032i | SRR3624698 |
| 053i | SRR3624700 |
| 117 | SRR3624701 |
| 12.1.014 | SRR14119027 |
| 12.1.037 | SRR14119026 |
| 12.1.078 | SRR14119025 |
| 12.1.181 | SRR14119024 |
| 12.1.205 | SRR14119023 |
| 12.1.217 | SRR14119022 |
| 12.1.225 | SRR14119021 |
| 12.1.234 | SRR14119020 |
| 12.1.241 | SRR14119019 |
| 12.1.291 | SRR14119018 |
| 127 | SRR3624702 |
| 169 | SRR3624703 |
| 204 | SRR3624705 |
| 205 | SRR3624706 |
| 37 | SRR3624699 |
| BTGP-6C4 | SRR14813096 |
| BTMaU-10 | SRR14813097 |
| BTMP-1828-5a | SRR14813098 |
| BTMP-1845-3 | SRR14813099 |
| BTMP-194-2 | SRR14813100 |
| BTMP-202-1 | SRR14813102 |
| BTMP-2807-4 | SRR14813101 |
| B71-2 | SRR6232156 |
| ZMW18_06 | ERR5525603 |
| ZMW18_08 | ERR5525605 |
| ZMW18_10 | ERR5525607 |
| ZMW18_11 | ERR5525608 |
| ZMW19_09 | ERR5525606 |
| ZMW19_13 | ERR5525609 |
| ZMW19_17 | ERR5525613 |
| ZMW20_03 | ERR5525601 |
| ZMW20_04 | ERR5525602 |
| ZMW20_07 | ERR5525604 |
| ZMW20_14 | ERR5525610 |
| ZMW20_15 | ERR5525611 |
| ZMW20_16 | ERR5525612 |
